# Vitexin inhibited the invasion, metastasis, and progression of human melanoma cells by targeting STAT3 signaling pathway

**DOI:** 10.1101/2020.09.24.311233

**Authors:** WenHao Zhang, LiPing Zhou, Guo Liu

## Abstract

In human melanoma cells, resistance to conventional chemotherapy and radiotherapy and rapid metastasis give melanoma a remarkable feature of the most aggressive and lethal. The low response rate of melanoma to existing treatment modalities is a substantial threat to patients and researchers. It is crucial to identify new therapeutic agents for the fatal malignancy melanoma. Vitexin is a flavonoid compound in many traditional Chinese medicines that exhibits antioxidant, anti-inflammatory and anti-tumour activities in many cancer cells. In our study, we elucidated the inhibitory effects of vitexin on invasion and metastasis in human melanoma A375 and C8161 cells *in vitro*. After vitexin treatment for 24 h or 48 h, the invasive ability and migration of melanoma cells were decreased in a dose- and time-dependent manners. In western blot analysis, we verified that vitexin inhibited the expression levels of MMP-2, MMP-9, vimentin, Slug and Twist which are known as the regulators of protein degradation and promote various cell behaviours such as migration and invasion. To further investigate the target signal that may be influenced by vitexin, immunofluorescence assay was performed to observe STAT3 localization and western blot results showed that vitexin decreased the expression of the phosphorylation of kinases that inducing STAT3 activation. Accordingly, we provide inspiring insight into the basic inhibition mechanism of vitexin, which will soon be an issue due to its scientific potential for further development as a novel anti-tumour agent for the clinical therapy of human melanoma.

## Introduction

With a poor prognosis and a progressively increasing incidence, cutaneous melanoma is attracting considerable interest as the most aggressive and deadly type of skin cancer(1). As a malignant tumour, melanoma originates from pigmentary cells originally derived from the neuroectoderm and can occur throughout the skin, iris, and rectum(2). Additionally, it has been reported that the third most common cause of brain metastasis is melanoma, following in lung and breast cancer, and more than 75% of melanoma patients suffer brain metastases, which cause death in 95% of all cases(3). The survival rate in patients with metastatic melanoma for 5 years, who receive traditional therapy is consistently <10% (4). However, chemotherapy, a strong anti-tumour strategy, yet fails to alleviate melanoma that shows drug resistance. Chemical drugs kill normal cells and cause toxic side effects that are unbearable for patients. In addition to fatal side effects, no obvious benefit in survival time is achieved with immunotherapy and targeted inhibitors(5). Consequently, developing novel therapeutic drugs is an urgent need, and such drugs should exert better anticancer activity, less toxicity, and the ability to coordinate current chemotherapeutic drugs to enhance their antitumor efficacy against human melanoma. As described above, cells can acquire invasive and migratory properties; these transformed cells progress to become life-threatening cancers(6). Degradation of stromal extracellular matrix (ECM) and basement membranes is one crucial step for malignant cells to invade and metastasize. As the family members of zinc-dependent matrix-degrading enzymes, matrix metallopeptidases (MMPs)(7, 8), MMP-2 (Gelatinase A, 72 kDa) and MMP-9 (Gelatinase B, 92 kDa) are two well-known gelatinases, and MMP-2 contributes to tumour cell migration after being directly activated by binding to αvβ3 integrin(9, 10). It has been reported that the expression of MMP-9 is vitally related to the invasiveness of melanoma cancer; thus, blockade of MMP-9 inhibits melanoma cells invasion(11). Epithelial-mesenchymal transition (EMT) is another metastasis pathway activated in cancer cells that also contributes to cell invasion and migration in various cancers(12-14). Some related proteins, such as vimentin, Slug and Twist, have been found to work synergistically in the regulation of EMT(15-17); thus, in this study, we sought to address whether vitexin could suppress the expression of EMT-related proteins and MMPs to further determine the underlying mechanism of vitexin inhibition.

Meanwhile signal transducer and activator of transcription 3 (STAT3), as an oncogenic transcription factor, plays a major role in malignant transformation(18, 19). During the progression of oncogenesis STAT3 is responsible for the transmitting of signals, from plasma membrane to nucleus, which results in the alternated expression of target genes related to cell proliferation, inflammatory response, metastasis, and immune response(18). In the STAT3 signaling pathway, the receptor-associated Janus kinase (JAK) that regulating the phosphorylation of STAT3 initially modulates the activation of STAT3 signaling through the recognition of specific cytokine and receptor of the cell surface(20, 21). Previous studies have indicated that vitexin inhibits tumour survival and invasion by suppressing the STAT3 pathway in hepatocellular carcinoma cells(22), and arrests the cancer cell cycle at G2/M phase to exert a broad-spectrum cytotoxic effect(23). Besides, other Chinese medicines such as quercetin, shikonin and dioscin have been reported to exert anti-melanoma effects through inhibiting the STAT3 signaling pathway(24-26). Due to the therapeutic potential of STAT3 inhibitors for melanoma treatment, many early-phase clinical trials are giving promising results. In current study, two human melanoma cell lines A375 and A2058 were used to verify the involvement of STAT3 in the anti-tumor effects of vitexin.

The aromatic shrub *Vitex negundo L* (Verbenaceae) is abundant in Asian countries due to its anti-inflammatory effect, has been widely used as a Chinese folk medicine for the treatment of asthma, cough, and arthritis. Recently the effective constituent of *vitex negundo L* was found to be vitexin (apigenin-8-C-glucoside), a naturally derived flavonoid compound, which is called ‘Mujingsu’ in Chinese(27). Vitexin is an active compound of *vitex negundo L*, but it can also be concentrated from various medicinal plants(28-31). Accumulating evidence shows that vitexin has a wide range of pharmacological effects including antioxidant, anti-nociceptive, and anti-Alzheimer’s disease effects(32-35). In particular the anti-tumour effect, which has been verified in many types of cancer (glioblastoma, pancreatic cancer, hepatocellular carcinoma and breast cancer)(36-38), has recently attracted increased attention. However, the characteristics of vitexin have not been dealt with in depth. A neglected area in the field about how vitexin may affect cell motility and migration in cancer; moreover, the underlying mechanisms of the suppression of invasion and metastasis in vitexin treatment remain unclarified in human melanoma cells.

In summary, the primary aim of this paper is to provide empirical and theoretical evidence for the claim that vitexin can inhibit the invasion and metastasis of melanoma cells *in vitro* through the suppression of the related proteins MMP-2 and MMP-9, and briefly discuss the involvement of STAT3 signaling pathway in these effects.

## Materials and methods

### Chemicals and reagents

Vitexin (purity >98%), which is purified from the seeds of the Chinese herb *Vitex negundo*, was purchased from Shanghai Yuanye Bio-technology Co., Ltd. (Shanghai, China), dissolved in dimethylsulfoxide (DMSO) and stored at −4°C until analysis. Dulbecco’s modified Eagle’s medium (DMEM), fetal bovine serum (FBS) and dimethyl sulfoxide (DMSO) were purchased from Gibco; Thermo Fisher Scientific, Inc. (Waltham, MA, USA). Matrigel was purchased from BD Bioscience (Pasadena, CA). Antibodies against MMP-2 (cat. No. 6E3F8), MMP-9 (cat. No. ab38898), vimentin (cat. No. 5741), Slug (cat. No. 9585), Twist (cat. No. 46702), GAPDH (cat No.5174), STAT3(cat. No. ab68153), phospho-STAT3 (Tyr705) (cat. No. ab76315), β-actin (cat No. ab8226) were purchased from Cell Signaling Technology, Inc. (Danvers, MA, USA). Antibodies against phospho-Src (Tyr416) (cat. No. ab40660), Src (cat. No. ab109381), phospho-JAK1 (Tyr1022/1023) (cat. No. ab138005), JAK1(cat. No. ab133666), phospho-JAK2 (Tyr1007/1008) (cat. No. ab32101), and JAK2(cat. No. ab ab32101) were purchased from Cell signaling Technology (Beverly, MA).

### Cell lines and cell culture

The human melanoma A375 and C8161 cell lines were acquired from Peking Union Cell Resource Center (Beijing, China). The cells were cultured in DMEM supplemented with 1% penicillin-streptomycin and 10% FBS with 5% CO2 in a humidified atmosphere at 37°C. The treatment groups of A375 and C8161 cells were treated with various concentrations of vitexin (5, 10, 15, 20 and 25 μM) at 37°C for 24 or 48 h, whereas cells in the control group were treated with equivalent volumes of DMSO.

### Cell viability assay

The effect of vitexin on cell viability was confirmed with Cell Counting Kit-8 (CCK8; Dojindo Molecular Technologies, Inc., Kumamoto, Japan) assay. A375 and C8161 cells (100 μl) were plated in a 96-well plate at a density of 1×10^4^ cells/ml and treated with various concentrations of vitexin (5, 10, 15, 20 and 25 μM). After 24 or 48 h, 10 μL of CCK8 solution was added to each well for 2 h at 37°C, according to the manufacturer’s protocol. The absorbance at 450 nm was measured using the iMark microplate reader (Molecular Devices, Sunnyvale).

### Wound-healing migration assay

To observe the migration of A375 and C8161 cells, an *in vitro* wound-healing assay was performed, following vitexin treatment. Cells (1×10 ^6^ cells/ml) were seeded into a 6-well plate and incubated for 24 h until they reached full confluence. The centre of the cell monolayer was scraped with a sterile 100-μL pipette tip to create a straight gap of constant width as a “wound”. Then, the cells were exposed to various concentrations of vitexin (0, 5, 10 and 20 µM) or DMSO (10 µM). After 24 h, 4% paraformaldehyde was used to fixed the cells, wound closure was imaged using a fluorescence microscope (Olympus Optical Co., Ltd., Tokyo, Japan), and the migrated cells were counted manually.

### Transwell invasion assay

To assess the motility of A375 and C8161 melanoma cells treated with vitexin, this invasion assay was performed using a modified Boyden chamber coated with 50 µl of Matrigel. The cells were then suspended at a density of 5 × 10 ^4^ cells/chamber and treated with vitexin (10 µM) or DMSO (10 µM). Medium containing 2% FBS was added to the upper wells while medium containing 10% FBS was added to the lower chambers. Then, the plates were incubated for 24 h at 37°C. Next, 4% paraformaldehyde was used to fix the invaded cells on the bottom surface of the insert, and the cells were stained with 0.1% crystal violet at room temperature. Images were obtained by using an inverted microscope (Olympus). The number of invaded cells was counted manually.

### Western blot analysis

Cells were cultured in 6-well plates at a density of 5×10^5^ cells/ml and treated with different concentration of vitexin (0, 5, 10 and 20 μM) at 37°C. After 24 h, cells were lysed in ice-cold radio-immunoprecipitation assay buffer (Beyotime Biotechnology Co. Ltd., Shanghai, China) containing a protease and phosphatase inhibitor cocktail for 30 min on ice. After centrifugation at 13,000 x g for 15 min at 4°C, the supernatant was collected. The protein concentration was quantified with a Pierce bicinchoninic acid protein assay kit (Pierce; Thermo Fisher Scientific, Inc.). Subsequently, equivalent amounts of protein (40 µg) were loaded, separated, and transferred to a polyvinylidene difluoride membrane (EMD Millipore, Billerica, MA, USA) with 10% SDS-PAGE. The membranes were incubated with primary antibodies (1:1,000; MMP-2, MMP-9, vimentin, Slug, Twist, GAPDH, phospho-Src, Src, phospho-JAK1, JAK1, phospho-JAK2 and JAK2; 1:2000; β-actin) after blocking in 5% skim milk at room temperature for 1 h and then incubating at 4°C overnight. Afterwards, at room temperature, the membranes were washed and incubated with horseradish peroxidase-conjugated secondary antibody (1:5,000) for 1 h. The membranes were washed three times with wash buffer 10 min. Proteins expression was detected using an enhanced chemiluminescence kit (EMD Millipore).

### Immunocytochemistry for STAT3 localization

To observe the localization of STAT3, A375 cells were treated with vitexin for 24 h and fixed with 4% paraformaldehyde. Then the cells were permeabilized with 0.2% Triton X-100, and blocked with 5% BSA in PBS. After that, A375 cells were incubated overnight at 4°C with specific primary antibodies against STAT3(1:100; Danvers, MA, USA). After being washed for three times, cells were incubated for 1 h with secondary antibody labeled with Alexa 488-labeled goat anti-rabbit IgG (1:200; Molecular Probes, Eugene, OR) at 25°C in the dark. After counterstained with DAPI, images of the cell signal were acquired under a confocal microscope (Nikon, Tokyo, Japan). During the observation of the cell signal, DAPI and FITC were excited at 405 nm and 488 nm, and the fluorescence emission was detected at 461 nm and 519 nm with 488 laserline and 568 laserline, respectively.

### Statistical analysis

All data represent at least three independent experiments and are expressed as the means ± standard deviations. Experimental differences were examined using ANOVA and Student’s t-tests, as appropriate. All statistical analyses were performed using SPSS v19.0 software (IBM Corp., Armonk, NY, USA). Statistical significance is indicated by *p < 0.05.

## Results

### Vitexin inhibits A375 and C8161 cell viability

The inhibitory effect of vitexin on human melanoma cell viability was determined. A375 and C8161 melanoma cell viability was measured by a CCK8 assay after treatment with various concentrations of vitexin (5, 10, 15, 20 and 25 μM). At 24 h, as presented in Fig. 1, cell viability was severely affected by high concentrations compared to low concentration treatment. As the concentration doubled, the cell activity showed a significant decrease. A similar result was observed at 48 h. The IC50 values were 22.46 μM (24 h) and 16.85 μM (48 h) for A375 melanoma cells and 17.59 μM (24 h) and 12.26 μM (48 h) for C8161 melanoma cells. However, compared to A375 cells, the slope of the graph is steeper in C8161 cells, and the values changes substantially between the concentrations of 10 and 20 μM, which may indicate that C8161 cells are more sensitive to vitexin than A375 cells when treated at 10 μM. Moreover, in both A375 cells and C8161 cells, 48 h of treatment led to a more significant suppression of cell viability than 24 h. In summary, the inhibitory impact of vitexin on A375 and C8161melanoma cells was a time- and dose-dependent.

**Figure 1.**
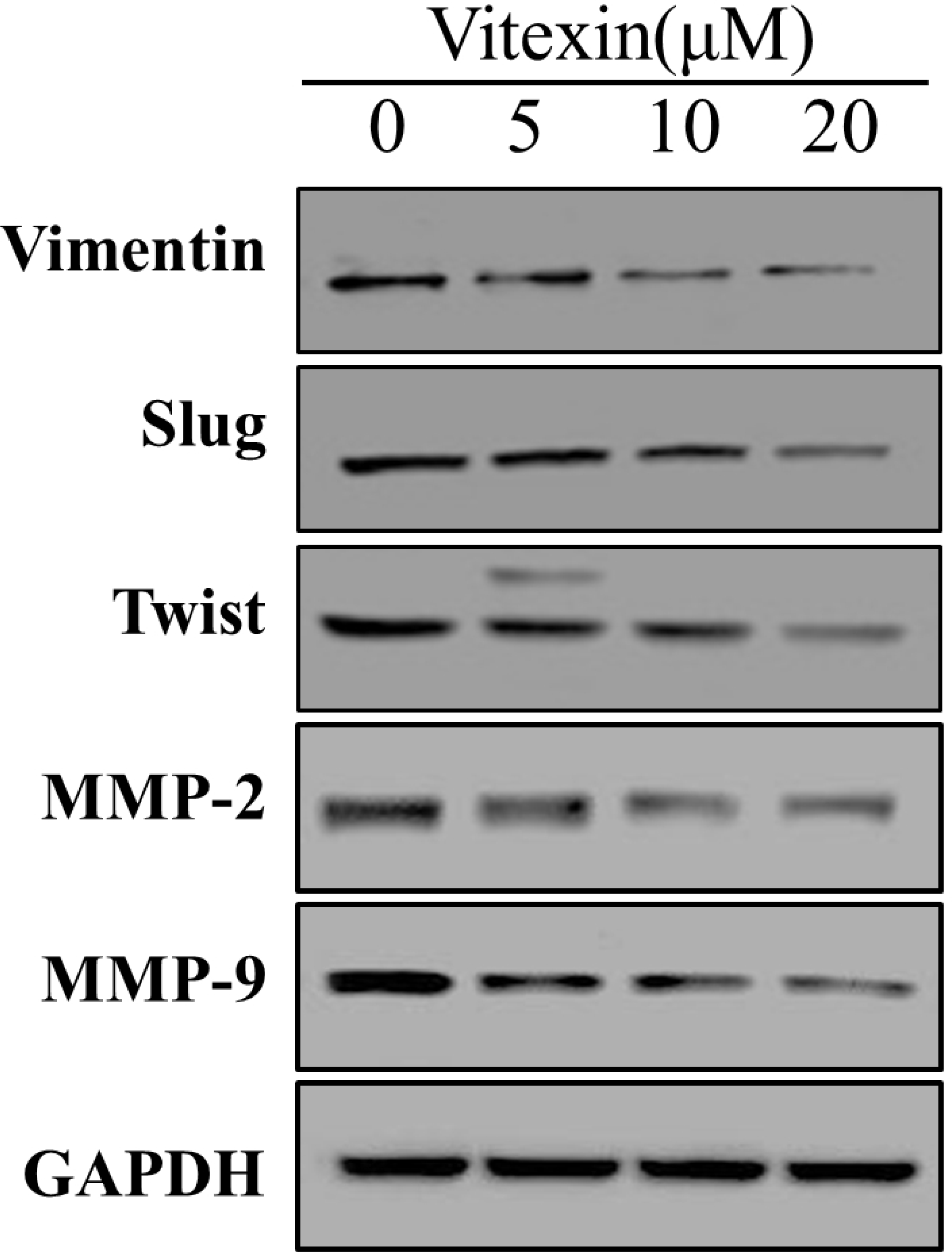
Inhibitory ability of vitexin on cell viability in the human melanoma A375 and C8161 cell lines. Cells were grown with various concentrations of vitexin (5, 10, 15, 20 and 25 μM) for 24 and 48 h. In addition, a CCK8 assay was done to assess melanoma cell viability. (A and B) The viability of A375 and C8161 cells was inhibited in a dose- and time-dependent manner after vitexin treatment. Data are presented as the mean ± standard deviation. * P<0.05, vs. control.

### Vitexin inhibits A375 and C8161 cell migration

A wound-healing assay was performed to investigate whether vitexin inhibits the migration of melanoma cells *in vitro.* A375 and C8161 cells were exposed to various concentrations of vitexin (0, 5, 10 and 20 µM) and the positive control (DMSO, 10 µM). At 24 h, the results revealed that vitexin reduced the migration of both A375 and C8161 cells in a dose-dependent manner (Fig. 2A,B and C).

**Figure 2.**
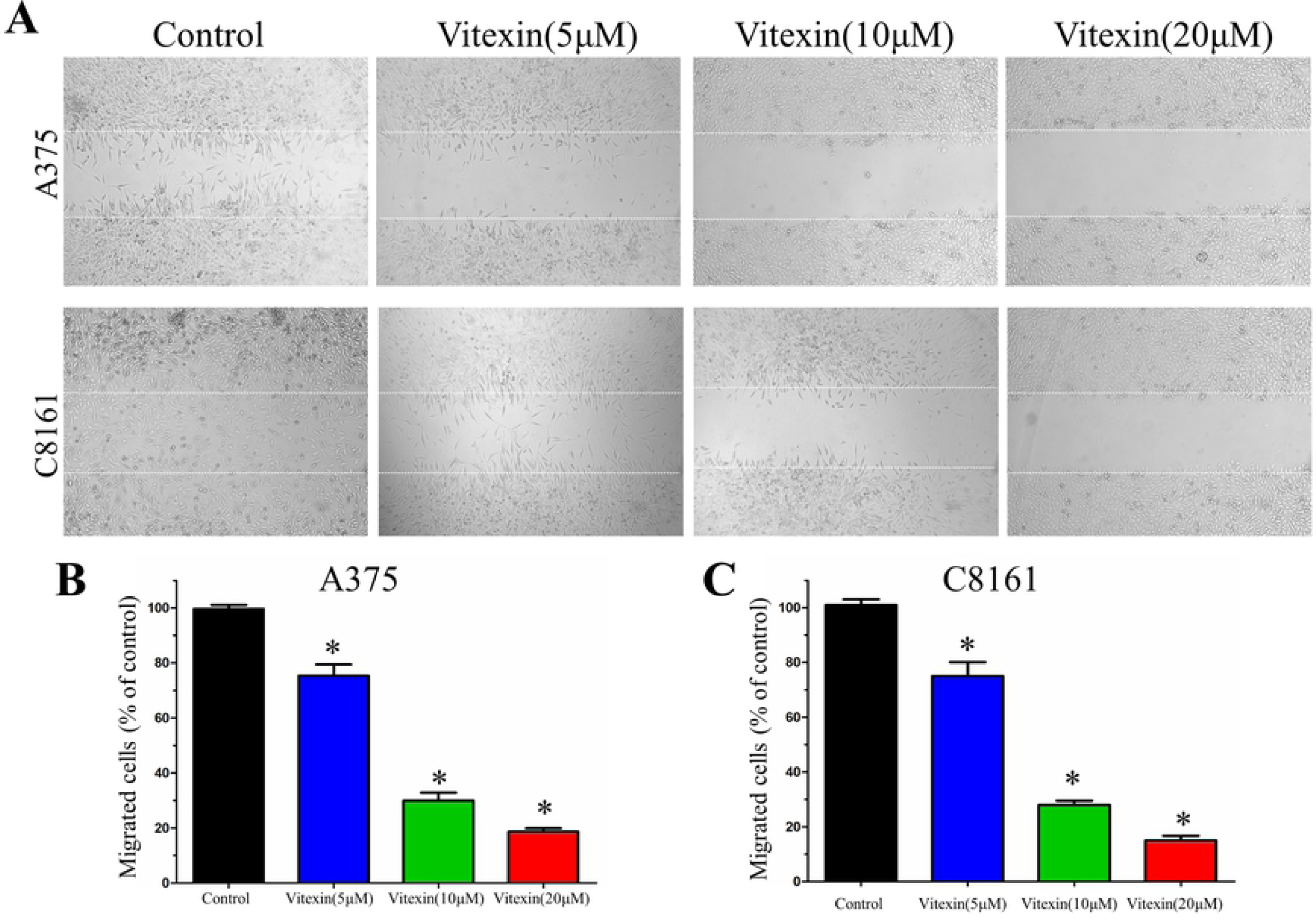
Vitexin inhibits A375 and C8161 cell migration. (A) In the wound-healing assay, vitexin inhibited the migration of A375 and C8161 cells in a dose-dependent manner. Scale bars = 100 μM. (B and C) The migrated A375 and C8161 cells were counted manually, and the degree of healing for each wound in the indicated groups is depicted in the histogram. Data are expressed as the percentage change (means ± SD). * P<0.05, vs. control.

### Vitexin inhibits A375 and C8161 cell invasion

In the Transwell assay, the invasion of A375 and C8161 cells treated with vitexin was observed. Following a 24 h treatment with vitexin (10 µM), there was a marked decrease in A375 and C8161 cells invasion, which indicated that vitexin can inhibit cell invasion at a low concentration in the Matrigel invasion assay (Fig. 3A). The number of invaded cells also demonstrated that when vitexin was added at a concentration of 10 µM, the inhibitory effect on cell invasion was almost the same in A375 and C8161 cells (Fig. 3B).

**Figure 3.**
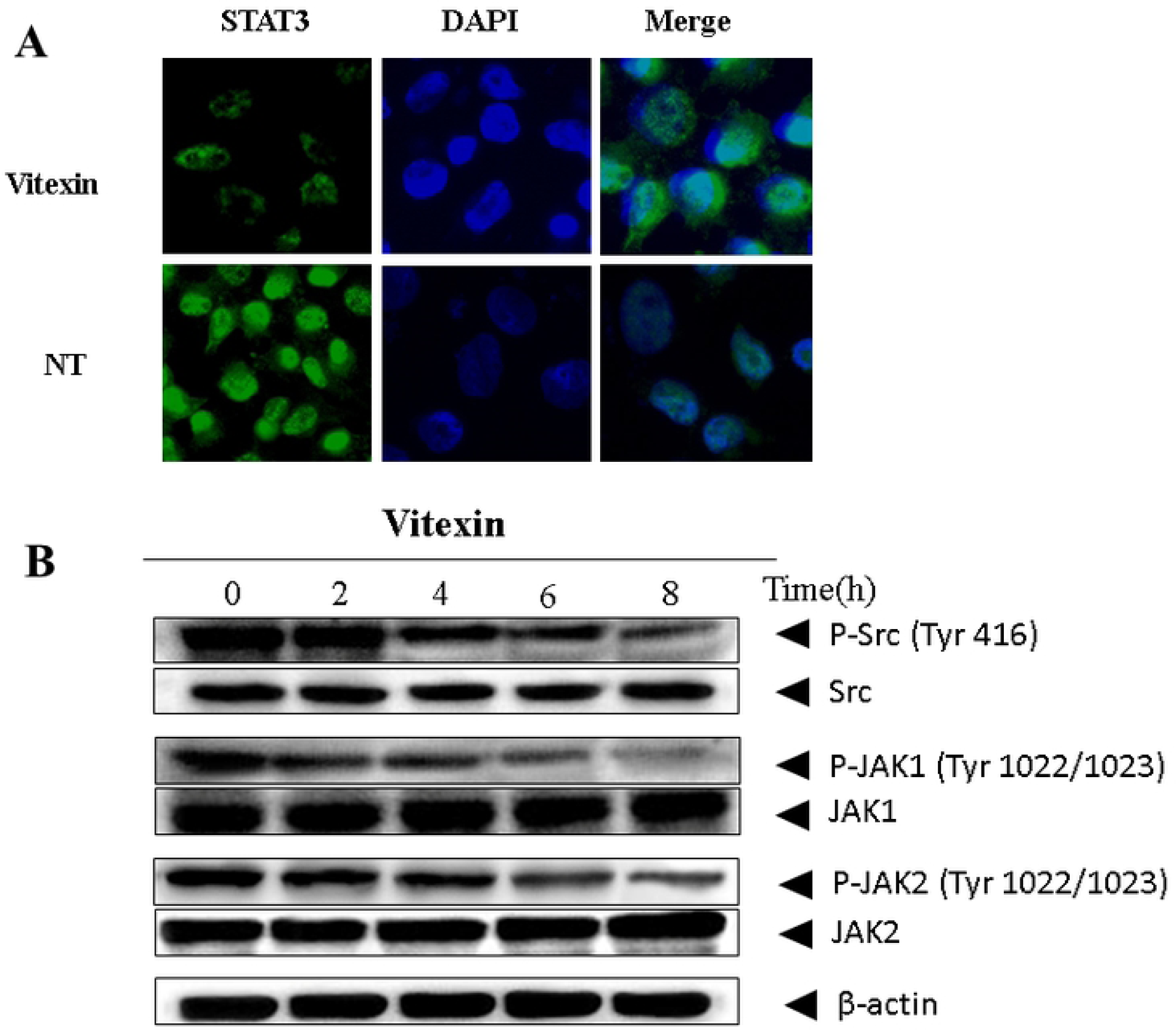
Vitexin inhibits A375 and C8161 cell invasion. (A) After 24 h of treatment with vitexin, cell invasion was evaluated by a Transwell invasion assay. Representative microscopic images of vitexin therapy are included. Scale bars = 100 μM. (B) Comparison of the rate of A375 and C8161 cell invasion. The relative proportions of invading cells with vitexin treatment are shown in the histogram. Data are expressed as the percentage change (means ± SD). P<0.05, vs. control.

### Vitexin suppresses the expression of migration-related proteins

To determine the proteins that vitexin may affect to inhibit cell migration, we performed western blotting on A375 and C8161 cells treated with vitexin (0, 5, 10 and 20 µM). Fig. 4 demonstrates that the expression of MMP-2, MMP-9, vimentin, Slug and Twist was reduced by vitexin treatment compared to no treatment. When melanoma cells were treatedwith different concentrations of vitexin (0, 5, 10 and 20 µM), the suppression of migration-related protein expression was dose-dependent.

**Figure 4.**
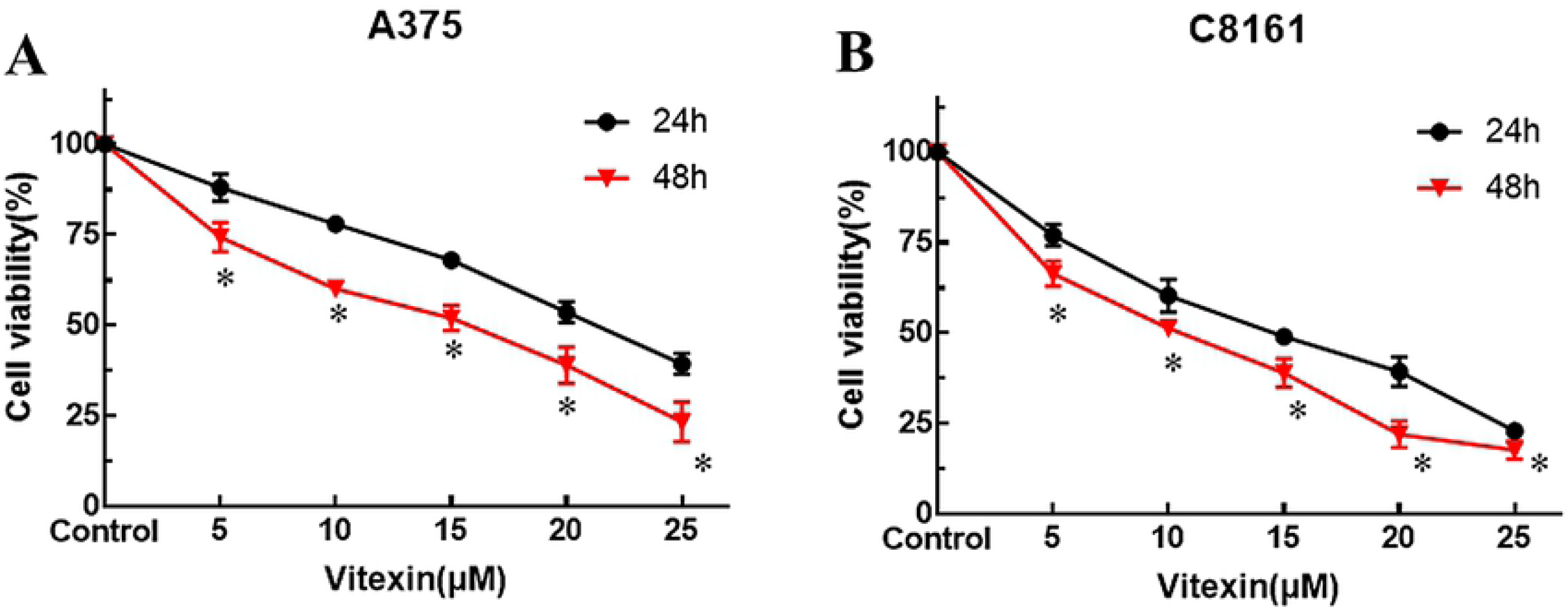
Vitexin suppressed the expression of migration-related proteins in a dose-dependent manner. A375 and C8161 cells were treated with different concentrations of vitexin for 24 h. The expression of MMP-2, MMP-9, vimentin, Slug and Twist were confirmed with a western blot assay.

### Vitexin inhibits the activation of STAT3 signaling pathway in melanoma cells

The cellular-distribution of activated STAT3 dimer plays a vital role for its transcription. Thus, STAT3 was determined by immunofluorescence assay to elucidate whether vitexin exposure can affect its nuclear translocation. Immunocytochemistry data indicated that treatment with vitexin downregulated the level of STAT3 in the nucleus in melanoma cells (Fig. 5A). To further explore the mechanism of vitexin on the inactivation of STAT3 signaling pathway, we performed western blot assay to evaluated the expression level of the phosphorylation of kinases that regulating the activation of STAT3. The regulation of STAT3 activation is closely associated with tyrosine kinases of the Src kinase and JAK families. As is shown in Fig. 5B, the expression of Scr and both JAK1 and JAK2 were persistently active. However, after vitexin treatment, the expression of phosphorylated Scr, JAK1 and JAK2 were significantly suppressed in a time-dependent manner (Fig. 5B).

**Figure 5.**
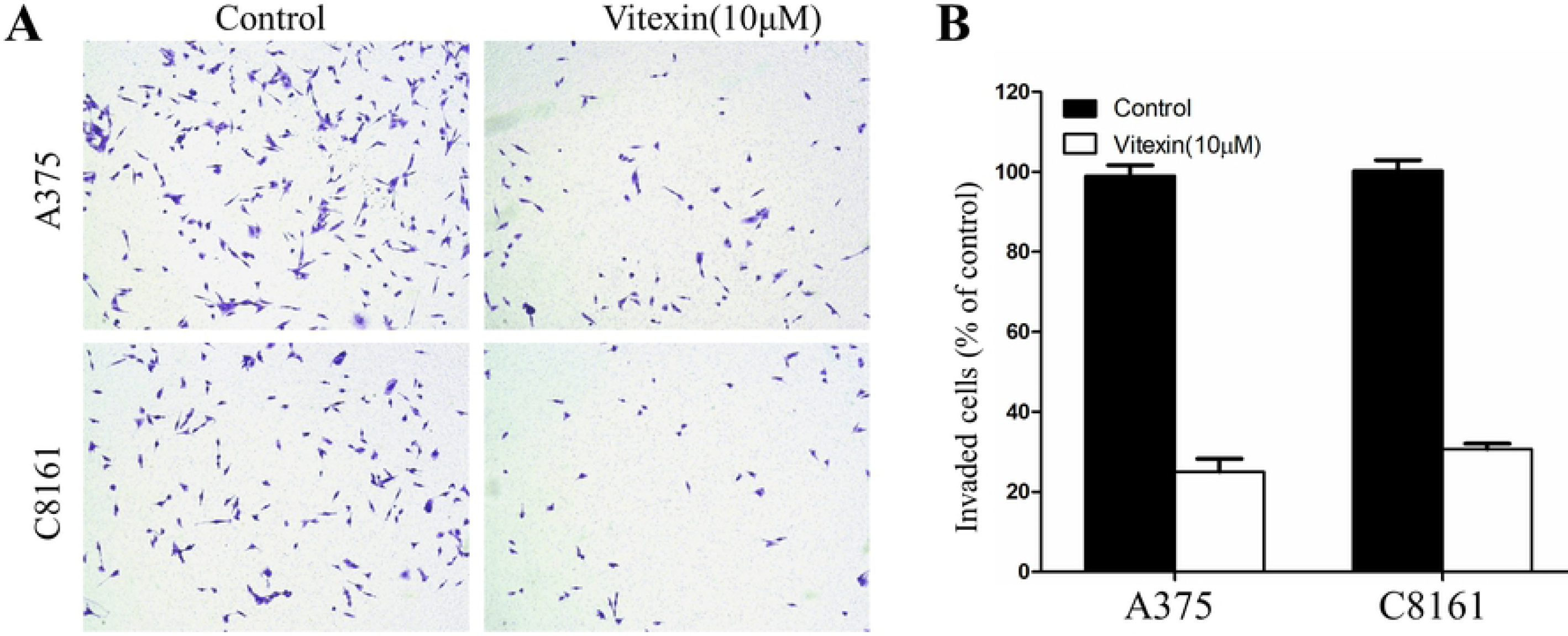
Vitexin negatively regulated the activation of STAT3 signaling pathway. (A) Human melanoma cell line A375 were exposed to vitexin for 24 h and vitexin altered the nuclear pool of STAT3 which was observed by immunocytochemistry. (B) A375 cells were treated with vitexin for the indicated time (0, 2, 4, 6, and 8h) and the expression of the phosphorylation of kinases regulating STAT3 activation were analyzed by western blot.

## Discussion

An epidemiological study has demonstrated that the major risk factors for developing melanoma are ultraviolet radiation exposure and severe sunburns(39). In the United States in 2016, melanoma contributed to 76,380 new cases and 10,130 cases of cancer-related mortality(40) and caused 75% of cases of skin cancer-related mortality. Global incidence of melanoma has increased to 15-25/100,000 individuals; melanoma-associated mortality also grows every year(41).Furthermore, metastatic melanoma tumours are frequently diagnosed with abnormal ectopic proliferation of melanocytes(42). Due to local invasion and migration, neither chemotherapy nor radiation therapy prolong length of life or increase living standards for patients with advanced melanoma(43). The spread of tumour cells from a primary site to remote sites is defined as metastasis, which is the final stage in tumour progression, representing a change from a normal cell to a fully malignant cell(44), and is the most common cause of death for cancer patients. Thus, developing effective anti-invasive agents as a therapeutic approach to control invasion and metastasis would be very likely to promote treatment.

Recently, vitexin, a plant-derived compound of *Vitex negundo L*, has attracted the attention of many researchers for its potential antitumor properties. Vitexin has a molecular formula of C21H20O10 and the chemical property is known as 8-D -glucosyl-4’,5,7-trihydroxy-flavone, or apigenin-8-C-glucoside(27). Pharmacologists revealed the quantity of stable radical (a total of 7) and the order of the hydroxyls in vitexin, 4′-OH > 7-OH > 5-OH, contributes to its bioactivities in cells(45). In particular, in the A ring of the vitexin structure, there was an o-di-hydroxyl structure that was verified to be responsible for effective radical scavenging in flavonoids(46). In terms of cell signaling, it has been reported that vitexin can affect different pathways, including p53, apoptosis and the cell cycle pathway, and arrest the cell cycle in G2/M phase by increasing the ROS levels in BRAFi (BRAF inhibitor)-resistant melanoma cells, which leads to DNA cytotoxicity and ultimately induces apoptosis in melanoma cells(29). Moreover, studies have revealed that there is strong potential for applying vitexin as a promising therapeutic Chinese medicine for cancer treatment such as non-small lung cancer and colorectal cancer, regardless of multi-drug resistance(47-49). Specifically, in PC12 cells(33), the anti-tumor metastasis effect of vitexin was verified, which inhibits HIF-1α (hypoxia inducible factor-1α) and reduces hypoxia-induced genes. The relationship between p53 and vitexin was also revealed in human oral cancer cells and indicated that via the p53-PAI1-MMP2 cascade, vitexin significantly suppressed cell viability and metastasis(34). However, the ability of vitexin to inhibit human melanoma cell invasion and migration has rarely been reported.

During tumour metastasis, the initial step is invasive tumour cell migration in the basement membrane and the proteolysis of the extracellular matrix, which involves several molecules, such as MMPs and integrins(50). Then, in invasive tumour cells, activated MMP-2 is recruited to the leading edge to cleave fibronectin into shorter fibronectin products, which can cleave various ECM proteins(7). Following activation of various intracellular signaling pathways, MMP-9 can be released by inflammatory cytokines, tumour necrosis factor-α, epidermal growth factor or phorbol ester (PMA)(51). In addition, transcription factors such as vimentin, Slug and Twist promote the EMT process that endows cancer cells with the abilities of invasion, migration and metastasis(14, 17). Therefore, agents acquiring the ability to suppress the expression of vimentin, Slug, Twist MMP-2 and MMP-9 should be developed to inhibit the invasion and metastasis of melanoma cancer. In the present study, data from *in vitro* experiments clearly demonstrated that vitexin is capable of inhibiting melanoma cell line viability in a CCK-8 assay. To determine the mechanism of vitexin in human melanoma cell inhibition, we also performed a western blot assay. As the image showed, the expression levels of MMP-2, MMP-9, vimentin, Slug and Twist are downregulated by vitexin in a dose-dependent manner.

Inspired by other studies, which have demonstrated that STAT3 could promote tumorigenesis in many malignant tumors by regulating several genes (52-54), the relationship between the anti-melanoma effects of vitexin and the involvement of STAT3 was elucidated in our study. As mentioned above, TWIST gene expression is responsible for cancer cell EMT which is known to be directly mediated by STAT3(55), and MMP-2 is also a STAT3-target gene that is involved in cell metastasis(56), indicating that inhibition of STAT3 is the possible mechanism of anti-migration in the anti-melanoma effects of vitexin. Through a reciprocal interaction with SH2 domain (a phosphotyrosine recognition domain), STAT3 is activated to be phosphorylated and transform to STAT3 dimers(57). Then, to transcribe various target genes, STAT3 dimers needs to translocate to the nucleus which was found to be inhibited after vitexin treatment in current study. Furthermore, it has been verified that STAT3 phosphorylation is regulated by upstream tyrosine kinases such as Src, JAKs and ABL(58). Therefore, we evaluated the effect of vitexin on Src, JAK1 and JAK2. Results showed that vitexin apparently mitigates the expression levels of p-Src, p-JAK1 and p-JAK2 in a time-dependent manner. Overall, these results demonstrated that in human melanoma cells, vitexin inhibits the expression of MMP-2, MMP-9, vimentin, Slug and Twist, which suppresses invasion and metastasis and exerts an adverse impact on tumour dissemination and metastasis. Moreover, the mechanisms of anti-melanoma effects of vitexin may partially attribute to the inactivation of STAT3 signaling pathway.

In conclusion, our present study revealed that vitexin could function as a novel inhibitor that decreased human melanoma A375 and C8161 cell lines migration and invasion by targeting STAT3 signalling pathway and suppressing the expression of MMP-2, MMP-9, vimentin, Slug and Twist. However, more experiments are required to thoroughly understand: (1) how vitexin downregulates the expression of matrix metallopeptidases; (2) whether vitexin suppress the expression of other STAT3 target genes or proteins, such as STAT3 driven oncogenic genes; (3) whether it is toxic for cancer patients subjected to vitexin treatment. Based on the results of current study, we provide further pharmacological groundwork of vitexin to be developed as a novel anti-tumour agent for the therapeutic strategies of human melanoma.

## Acknowledgements

This study was supported by the Science Foundation of Shandong Province (grant no. S20160921).The authors declare that they have no competing interests.

